# Efficient and effective identification of cancer neoantigens from tumor only RNA-seq

**DOI:** 10.1101/2024.08.08.607127

**Authors:** Danilo Tatoni, Mattia Dalsass, Giulia Brunelli, Guido Grandi, Mario Chiariello, Romina D’Aurizio

## Abstract

The growing accessibility of sequencing experiments has significantly accelerated the development of personalized immunotherapies based on the identification of cancer neoantigens. Still, the prediction of neoantigens involves lengthy and inefficient protocols, requiring simultaneous analysis of sequencing data from paired tumor/normal exomes and tumor transcriptome, often resulting in a low success rate. To date, the feasibility of adopting a more efficient strategy has not been fully evaluated. To this end, we developed ENEO, a computational approach to detect cancer neoantigens using solely the tumor RNA-seq data while addressing the lack of matched control through a Bayesian probabilistic model. ENEO was assessed on TESLA benchmark dataset, reporting efficient identification of DNA-alterations derived neoantigens and compelling results against state-of-art exome-based methods. We further validated the method on two independent cohorts, encompassing different tumor types and experimental procedures. Our work demonstrates that a tumor-only RNA-based approach, such as the one implemented in ENEO, maintains accuracy in identifying mutated peptides resulting from expressed genomic alterations, while also broadening the pool of potential pMHCs with RNA-specific mutations in a faster and cost-effective way. ENEO is freely available at https://github.com/ctglab/ENEO

## Introduction

Somatic mutations responsible for the change of the aminoacidic sequence of a given protein are referred to as non-synonymous, and could lead to the generation of tumorspecific neoantigens (1). The absence of these mutations from the healthy human genome makes these peptides not subjected to self-tolerance, increasing the attractiveness towards their use for the development of targeted immunotherapies (2).

Owing to the growing accessibility of high-throughput sequencing experiments, the development of individualized immunotherapies based on patient-specific neoantigens has been significantly sped up, showing promising results in multiple cancer types, such as melanoma (3, 4), gastrointestinal cancers (5, 6) and even particularly hard-to-target brain tumors as glioblastoma (7). The engineering of these therapies requires the identification of cancer neoantigens with a predicted high binding affinity towards the patient specific Human Leukocyte Antigen (HLA) (1, 8). The identification of patient specific neoantigens is a critical step which typically starts by calling somatic variants using genomic data, from either the whole exome (WES) or whole genome (WGS) sequencing, obtained from the patient’s tumor and healthy tissue (8). The expression of the resulting aberrant transcripts must be then confirmed, requiring the concurrent profiling of tumor RNA-seq data (9). The obtained mutations are then used to generate candidate peptides whose binding affinity towards major histocompatibility complex (MHC) is predicted using computational tools (8).

Although widely adopted, this approach is not exempt of limitations. Even though the genomic profiling of cancer mutations could results in a notable set of predicted candidate neoantigens, only a minimal fraction of them may result in the effective stimulation of an immunogenic response in the patient (5, 6, 10). Indeed, while WES opens to the high throughput profiling of nearly all the open reading frames of the tumor genome, DNA variants falling in un-transcribed regions is not predicted to generate neoantigens (1). Moreover, with the increasing adoption and optimization of protocols for profiling the MHC-I restricted immunopeptidome in cancer (11), it has become clear that DNA variants alone failed to fully explain the variety of detected peptides (12–14). Conversely, the profiling of RNA-alterations using RNA-seq from the same cell lines (12, 14) or from the same tumor biopsy (14–16) empowered the spectra resolution, demonstrating how alterations detectable from tumor RNA offer a consistently better representation of the MHC-presented peptides space. These evidences were supported and enforced by the recent work of Tretter and colleagues (17), which demonstrated, using a wide cohort of patients spanning multiple tumor types, that only a small fraction of immunogenic neoantigen candidates detected by mass spectrometry could be explained using DNA variants.

Detecting somatic alterations from the tumor RNA-seq alone has been proven to represent a feasible approach (18), even though not exempt of troubles (19). Due to the nature of the RNA-seq experiment, selecting an appropriate matched control remains complex and not fully resolved (20). Additionally, RNA-seq reads are more error-prone due to the splicing mechanisms, which may result in alignment errors nearby splicing junctions and reverse transcriptase artifacts (21). Despite these obstacles, advancements in variant calling algorithms have driven the exploration of tumor-only RNAseq methods for capturing tumor mutational burden and identifying driver mutations across various cancer types (18).

To date, the feasibility of adopting solely the tumor RNAseq for the effective prediction of immunogenic neoantigens in a personalized immunotherapy scenario has not been fully evaluated. To this end, we developed ENEO, an easy-to-use computational workflow encompassing all necessary analytical steps, from variant calling to pMHC binding affinity, in a reproducible and scalable manner. ENEO addresses the absence of a matched control sample by utilizing a Bayesian probabilistic model that takes advantage of genetic population databases. We demonstrate that ENEO effectively detects tumor neoantigens across various tumor types and experimental setups using publicly available data. These analyses open to the use of tumor RNA-seq as a rapid and costeffective method for detecting tumor neoepitopes, enhancing the role of deep transcriptional profiling in precision oncology.

## Materials and Methods

### Data used in this study

We downloaded matched WES data (tumor/normal) and tumor RNA data of five patients generated by the TESLA consortium (22), 3 of which with melanoma (namely TESLA_1, TESLA_2, TESLA_3), and 2 with non-small cell lung cancer (TESLA_12, TESLA_16) from Synapse (syn21048999, access granted). Tested neoepitopes and their relative validation response were collected from the supplementary materials attached to the related publication (22). RNA-seq data of the 3 melanoma samples from Gros et al (23) were downloaded from SRA (SRP064661) and 8 patients with gastric cancer from Tran et al (5) and Parkhurst et al. (6) from SRA (SRP278662). Within the last cohort, the selected patients are those with a single sequencing experiment carried out from the resected metastasis. Peptide validation results for these two cohorts, together with the detected somatic variants, were obtained from the retrospective dataset built by Gartner et al. (24).

### Identification of germline and somatic variants from WES

WES data were preprocessed using *fastp* (25) (v.0.23.2) to remove sequencing adapter and low quality reads. Reads mapping was performed using *bwa* (26) (v.0.7.17) with default parameters and the GRChg38 human genome assembly as reference. Resulting alignment files were then processed following the GATK Best Practices (27). Somatic variants were called from paired tumor/matched control with *Mutect2* (28) (v.4.2.0) and subsequently filtered as previously described (27). Germline variants were called from control WES using *DeepVariant* (29) (v.1.6.0) using default parameters. To exclude variants falling in known challenging regions of the human genome (30, 31), we downloaded target lists from the Genome In A Bottle (GIAB) ftp^1^ and removed overlapping variants using *bedtools* (32) (v.2.30.0). Variant effect prediction annotation was performed on both the call sets using VEP (33) with the Ensembl version 105.

### Identification of candidate neoantigens from tumor RNAseq with ENEO

We implemented a modular and reproducible computational pipeline within the Snakemake framework (34), named ENEO, to identify putative neoantigens using solely tumor RNAseq data (depicted in **Figure 1**). The overall workflow encompasses three main step which are described in full detail below.

**Fig. 1.**
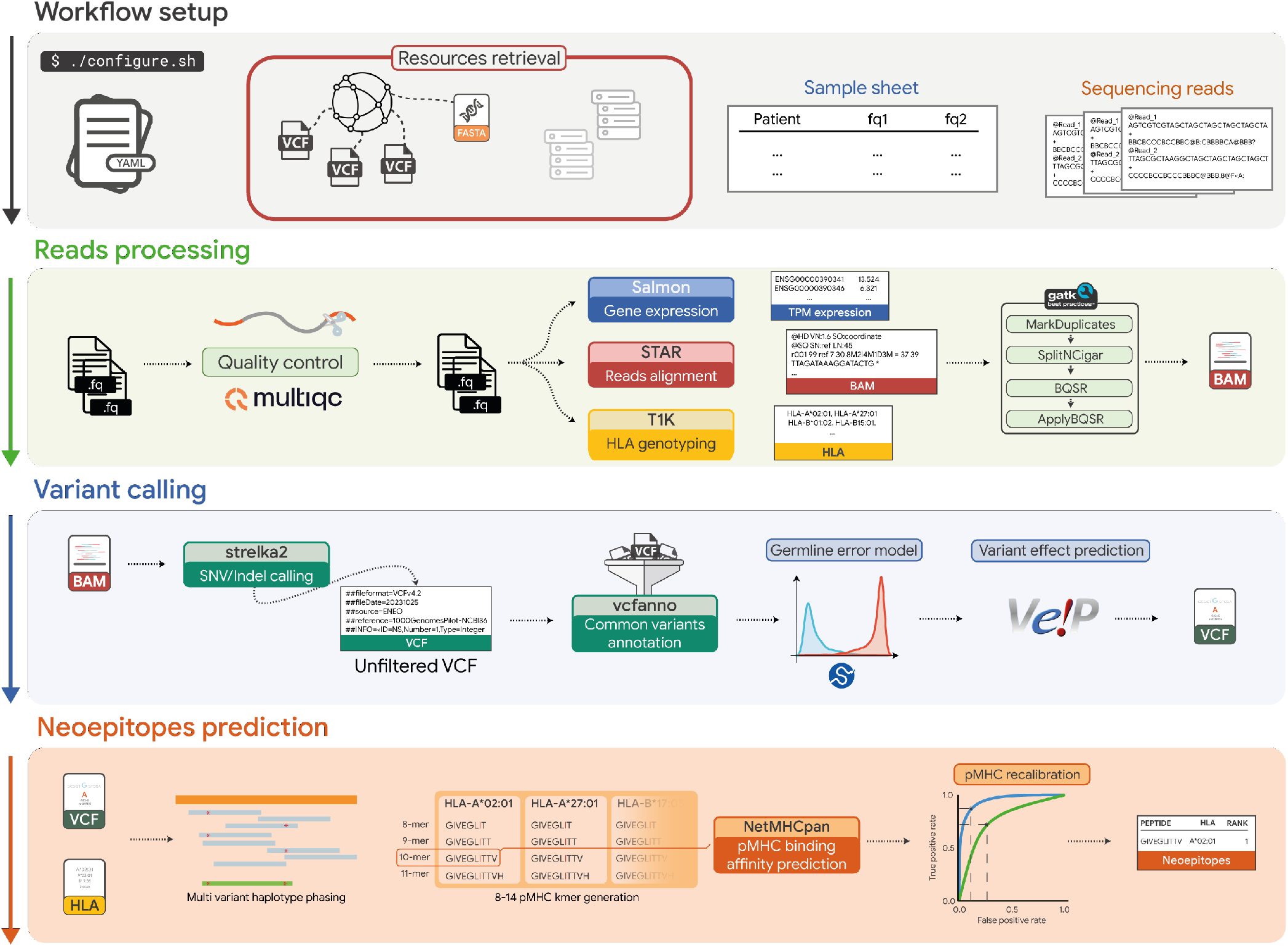
Overview of the ENEO computational workflow. The pipeline is composed of three main phases: Reads Processing (1^st^ step), Variant Calling (2^nd^ step) and Neoepitopes Prediction (3^rd^ step). Input files required by the workflow are indicated on the top grey box, together with the employed public resources whose first download and setup is guided. Intermediate post-processed output files (i.e. BAM, VCF, HLA) used throughout the pipeline are saved and made available for further user inspection.

### Reads alignment and file processing

First, tumor paired end RNA-seq data are preprocessed to trim adapter sequences and low-quality reads using *fastp* (25) (v.0.23.2). To increase the read confidence, the base correction method of *fastp* is adopted. It corrects poor-quality bases (PHRED < 14) in a read if the other sequencing pair has a discordant base with excellent sequencing quality (PHRED > 30), given an overlapping region with a minimum length of 30 bases. Reads are then aligned with STAR (35), using the GRChg38 assembly genome and the Ensembl 105 annotation, with the “*2Pass-mode*”, to increase specificity by generating sample-specific exon-splicing junctions and to enhance the overall mapping confidence. Aligned reads are further processed according to the GATK Best Practices (27) for short variants discovery from RNAseq. In short, duplicate reads are flagged using *MarkDuplicates* (v.4.2.0) to discard PCR and optical duplicates; aligned reads are then passed through *SplitNCigar* (v.4.2.0), that is responsible for the splitting of those containing ‘N’ in their CIGAR string and for the hard-clipping of overhanging regions containing mismatches (a common scenario across exon junctions); the base quality is then recalibrated using Base Quality Score Recalibrator (BQSR) (v.4.2.0) to adapt for the subsequent variant calling step.

### Variant calling, filtering and annotation

The variant calling process is performed with *strelka2* (36), using the argument “*rna*”. Despite being designed originally for germline variant calling, its relaxed modeling of genotypes enables the inclusion of the plethora of different measured allele frequencies coming from somatic alterations of different clonality (17, 36). Variants are called over a provided set of all protein coding exons, as obtained from the Ensembl v.105, after the exclusion of the genes belonging to the antigen processing and presentation. The list of excluded genes and their relative chromosomal regions is reported in the **Supplementary Table 1**. To ensure a minimum evidence boundary for candidate somatic variants, *bcftools* (37)(v.1.19) is used to remove alterations in regions not covered by at least 5 reads and those with fewer than 3 reads spanning the alternative allele. Similar to the WES case, and based on the same motivations to select only high-quality variants, those detected in challenging regions of the human transcriptome are removed. To report common germline events, candidate variants are annotated with *vcfanno* (38) (v.0.3.5) using multiple population databases: exAC (39)(r. 1), gnomAD (40) (v.3.1) and the ALFA project (41). The obtained allele counts and frequencies are weighted using geometric average, to account for the different sample size in each dataset. Variants’ consequence annotation is performed with VEP (version 105), using additionally the plugins *Wildtype* and *Frameshift* to report the protein sequence within the VCF file.

### Definition of a Bayesian probabilistic germline error model

As the variant calling process of *strelka2* already accounts for potential systematic errors of the RNA-seq experimental setup, we attempt to account for the confounding effect caused by the simultaneous presence of both germline and somatic alterations in the pool of variants called from tumor-only transcriptomic data. To this end, ENEO employs a Bayesian probabilistic model, which derives the likelihood of a called variant being germline in the diploid scenario. The model leverages, when available, the measured population allele frequency retrieved from the germline resources (i.e. exAC, gnomAD and ALFA project), here denoted as *f*. In contrast, whenever *f* is not known, we adopted the same approximation introduced by Benjamin and colleagues in the Mutect2 paper (28). Briefly, the number of variant alleles n in a germline resource composed by *K* patients, with *N* = 2*K* chromosomes, could be modeled as coming from a binomial distribution as *n* ∼ *Binom*(*N, f*). Supposing a prior for the frequency *f* to be *f* ∼ *Beta*(*α,β*), centered on the average human heterozygosity 𝔼 (*f*) ≈ 10(− 3), the probability of sampling a site not found in the germline resource is the result of a beta-binomial process as *P* (*n* = 0) = *BetaBinom*(0|*α, β, N*). This could be approximated to the measurable number of exonic sites not mutated, resulting in *P* (*n* = 0) ≈ 7*/*8. Updating the Beta prior on *f*, it results in a posterior *f ∼ Beta*(*α, β* + *N*) whose mean value is approximated to *α/N*, given that *N* ≫ *β*. Using the gnomAD 3.1 reference (*N* = 71702) we obtained a mean value approximal to 7*e*^*−*8^ which is used as *f* for variants not found in the germline resource.

Considering the diploid scenario, the unnormalized probability of three main events is defined as follow:

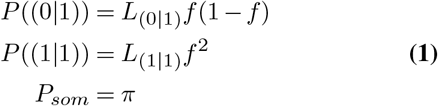

Where:

- *L*_(0|1)_ and *L*_(1|1)_ represents the likelihood emitted by *strelka2* for the given variant, respectively, for the heterozygous and homozygous genotype
- *f* is the population allele frequency derived from the population databases as previously described
- *π* is the somatic occurrence ratio

The parameter *π* inside the probability estimation represents the likelihood of observing a somatic event in a position within an exon of a protein coding gene, without accounting for the given clonality and/or ploidy. We modeled it as the results of two subsequent Bernoulli processes, where the first represents the successful event of finding an exon *E* that contains at least a variant, and the second as the successful event of finding a mutated base *B* out of all the bases *n* of that exon. Considering each exon *E* as independent each other, finding a mutated exon *E*_*MUT*_ is dependent upon the probability *p*_*E*_ and could be defined as:

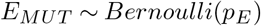

Within an altered exon *E*_*MUT*_, considering each of the composing bases as independent each other, the probability *p*_*B*_ of picking the position borrowing the variant is defined as the conditioned probability

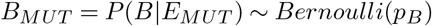

This entangles the model as dependent on two distinct probabilities which, to encompass the different mutation susceptibility observable throughout the human genome, could not be coerced to fixed values. We then suppose an independent prior Beta distribution for each, defined as:

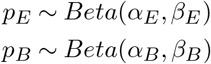

To estimate the parameters of these two Beta distributions, we collected somatic variants of 2683 cancer patients from the PCAWG consortium through cBioPortal (cbioportal.org). We retained only those alterations falling within the regions used by ENEO for variant calling to ensure consistency. To speed-up model convergence, we excluded patients with fewer than 50 somatic calls, resulting in a final set of 1523 patient across 26 tumor types. From each patient we collected the observed probability of a mutated exon 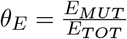 and the observed probability of a mutated base 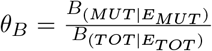 for all the exons. Given the fact that both the parameters of a Beta distribution must fall in ℝ^+^, we assigned non-informative Gamma distribution as priors

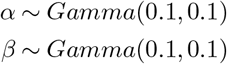

Non-informative priors were chosen to do not enforce beliefs about the parameter values, allowing the data to inform the posterior estimates. We employed Metropolis Hastings algorithm (42) to approximate the posterior distributions of *α* and *β* based on the collected data. The algorithm was run for 5000 iterations with a burn-in period of 1000 iterations to ensure convergence and reduce the influence of initial values. The highest probability density estimated for each parameter was then used to update the respective parameters of both the Beta distributions. This approach allows us to identify the most likely parameter values given the observed data. Model inference was conducted using the *scipy* (43) library (v.1.12) in Python (v.3.10).

### Gene expression quantification and HLA typing

Transcript expression quantification is conducted using *salmon*(44) (v.1.9.0), and subsequently summarized at genelevel TPM (transcript per million) using the R package *tximport*(45)(v.1.30). HLA genotyping is performed with T1K (46) (v.1.0.1) using raw reads (FASTQ) as input. Only the genotypes for the HLA-(A|B|C) loci with the highest coverage and likelihood are kept as input for the latter stage.

### Prediction of neoantigens binding affinity

Annotated variants and patient HLA genotypes are fed for the last step of ENEO, which is responsible for the generation of candidate neoantigens and the prediction of their binding affinity. To increase the specificity of predictions, we used the haplotype phasing information to obtain multivariant phased representations of the altered protein, as reported previously (47). From the resulting mutated aminoacidic sequence, all k-mer peptides (from 8 to 14aa) which include the mutated position are extracted moving a sliding window through the sequence. These peptides, along with each of the patient’s HLA alleles, are then passed to the NetMHCpan 4.1 (48) algorithm, which predicts the binding affinity of the peptide to the HLA molecules. This results in a list of pMHC complexes, with the binding likelihood expressed as a percentile rank against a set of random peptides. To obtain an high confidence value specific for a given allele, instead of adopting a fixed acceptance boundary for the percentile rank, we resembled the approach adopted by Reardon and colleagues (49). After obtaining pMHC data from the IEDB (50) v3 (iedb.org) and training data of NetMHCpan4.1 from the supplementary materials of the publication (48), we generated the negative background peptide dataset by randomly slicing k-mer peptides within the length range of 8-14 from the human RefSeq proteome (GCF_000001405.40). Then, after removing pMHC pairs already present in the training data, we kept positive and negative pairs until reaching a proportion of positive:negative of 1:9. To ensure minimum confidence, we proceeded using only alleles with at least 100 experimentally validated unique peptides. For each allele, we used the peptides’ predicted percentile ranks by NetMHCpan 4.1 and the corresponding labels to obtain the optimal percentile threshold, defined as the value that maximizes the difference between the true positive rate (sensitivity) and false positive rate (1 specificity). This resulted in the identification of HLA-specific optimal values for 114 different HLA alleles, which are used to filter the set of candidates pMHC arising from the detected variants. Natural non-malignant HLA ligands, potentially resulting from healthy tissues and so subjected to self tolerance, are removed using the data collected from the HLA Ligand Atlas (51) (release 2020.12). Resulting neoantigens are then ranked according to their predicted percentile rank and reported in the ENEO output along with the expression of the mutated gene, opening for optionally additional ranking strategies.

## Results

### Comprehensive identification of cancer neoantigens from tumor-only RNA-seq with ENEO

To identify and prioritize putative candidate neoepitopes using solely the tumor transcriptome, we developed ENEO, a scalable, reproducible and extensible computational workflow implemented using the Snakemake (34) framework. ENEO’s workflow encompasses three main steps (i.e. “Reads processing”, “Variant calling” and “Neoepitopes prediction”), whose graphical representation is depicted in **Figure 1** and detailed in the **Methods** section. The first step is responsible for the reads pre-processing, mapping and alignment post-processing, accomplishing three main tasks: the gene expression quantification, the delivery of a processed alignment file compliant with GATK Best Practices, and the HLA genotyping. The second step implements strelka2 for variant calling, the ENEO’s germline error model for somatic variants likelihood estimation and the functional annotation with VEP.

In the absence of a matched normal sample, discriminating between germline and somatic variants is crucial. Here, we present a Bayesian probabilistic model that derives the likelihood of observing a somatic event using either the calculated or estimated germline population frequency and the genotype likelihood derived from sequencing data. In our model, we assumed that the likelihood of identifying a somatic variant within a specific coding exon is contingent upon two Bernoulli trials. The first involves sampling a mutated exon out of the pool of expressed protein-coding exons, while the second regards sampling the mutated nucleotide. We assumed that the probabilities of both events follow a Beta distribution, characterized by two shape parameters (i.e. *α* and *β*). Using data from 1523 patients across 26 tumor types from the ICGC (see Methods), we approximated the posterior distributions for these parameters using the Metropolis Hastings algorithm. For the exon mutation probability, the posterior mode for *α* and *β* were found to be 0.87 ([0.78,0.96] highest probability density (HPD) interval) and 612.23 ([532.92,702.62] HPD interval) respectively. Conversely, we obtained 0.95 ([0.94, 0.96] HPD interval) and 199.58 ([197.98, 203.61] HPD interval) as the posterior mode for *α* and *β* of the probability of sampling the mutated nucleotide, The posterior density plot confirms that our model proposes for both a highly skewed Beta distribution, that fits accurately the observed data distribution (**Fig. S1**). Probabilities sampled from the target posterior distributions are used, together with the population allele frequency and the variant genotypes’ likelihood (Methods), to annotate variants in the resulting VCF file with the estimated probabilities. The output of the second step is a VCF file with the full set of variants along with their variant effect prediction, used to determine the resulting mutated protein sequences. Lastly, in the third step, all variants are then used for generating candidate peptides and, coupling with the HLA genotype, the pMHC binding affinity is then predicted using NetMHCpan 4.1. The final output of ENEO is a ranked list of patient’s specific candidate neoepitopes arising from somatic mutations. ENEO is freely available at https://github.com/ctglab/ENEO.

### Efficient disjoin of germline variants from the tumor-only RNA callset through ENEO germline error model

We evaluated the ability of our model to identify most of the germline variants present in tumor cells, given the absence of the genetic profile of matched non-tumor cells, by comparison with the standard calling approach which exploits both the DNA sequencing of tumor and paired non-tumor samples (T/N WES). To accomplish that, we used data provided by the TESLA consortium (22). These data, indeed, include exome and transcriptome sequencing of cancer biopsies collected from 5 patients with melanoma and non-small cell lung cancer, along with exome sequencing from PBMCs as healthy controls (**Methods**).

At first glance, we tested the sensitivity of the implemented germline error model to correctly distinguish the provenience of reported variants, either germline or somatic. To do so, we called germline alterations from control WES using DeepVariant (**Methods**) and somatic alterations from T/N WES using Mutect2 (**Methods**). Despite the different algorithms employed, no overlap was observed between germline and somatic WES calls across all patients (**Fig. S2**). We obtained on average 4,343 somatic WES-variants with predicted effect on downstream protein, with melanoma patients showing an overall higher number of variants (**Fig. S2**), in line with the expected burden of the tumor types under investigation (52). Within the intersection between WES-variants and RNA-variants from ENEO, we assessed the sensitivity of the germline error model. Starting from a set of 23,762 RNA-variants detected also from germline WES analysis, the adoption of a probability cutoff of 0.5 resulted in the successfully labeling of 23,296 (∼ 98%) variants, showing consistent performance for all the patient under investigation (**Fig. 2A**).

**Fig. 2.**
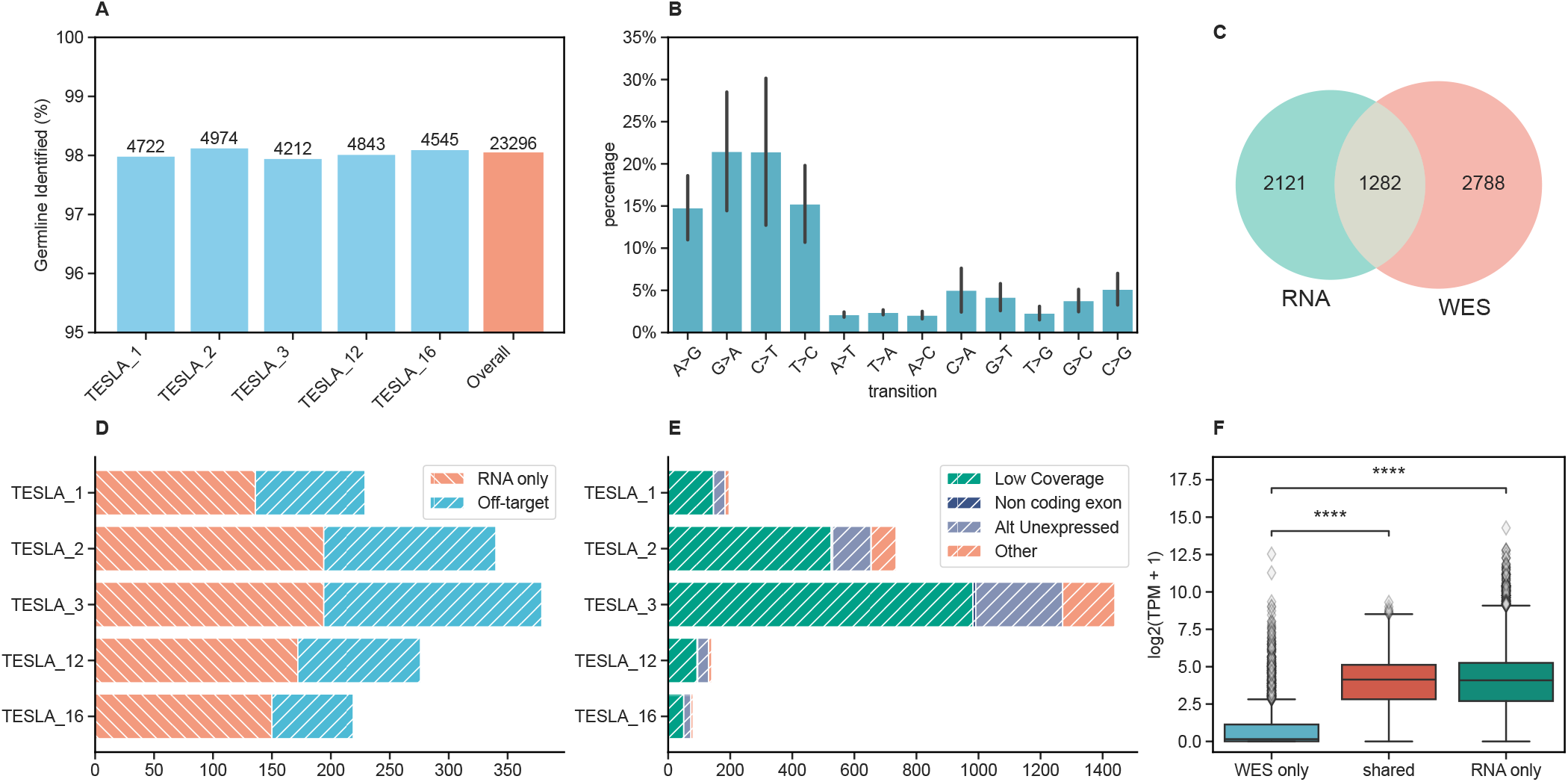
Characterization of somatic variants identified by ENEO in comparison with variants identified from WES in the TESLA cohort. **A**. Percentage amount of germline variants in patients of the TESLA cohort correctly identified by the germline error model of ENEO. **B**. Distribution of ENEO candidate somatic variants within each single nucleotide variant type. Percentage data from each patient are aggregated in the same bar. **C**. Overlap between the candidate calls identified by ENEO “RNA” with somatic alterations identified from tumor/normal variant calling “WES” (see *Methods*). **D**. Distribution of variant unidentified from WES divided in: “Off-target” (refers to variant occurring in a region out of the exome capture kit) and “RNA only” (refers to polymorphism only detected from ENEO). **E**. Distribution of variant unidentified by ENEO divided in: “Low Coverage” (refers to a locus with overall coverage below the calling threshold of ENEO, see Methods), “Non coding exon” (describes regions outside of the target regions employed by ENEO), “Alt Unexpressed” (refers to regions that, despite being transcribed, failed to show evidences of alternative allele expression) and “Other” (collects cases not covered by the other categories). **F**. Expression level of genes reporting variants only detectable from WES, (“Wes only”), only from RNA (“RNA only”) or in both assays (“Shared”). Expression measured using transcript per million (TPM). Kruskal Wallis test. **** p ≤ 0.0001

After filtering, we retained a total of 3,403 candidate somatic variants with a predicted impact on the downstream protein sequence across all patients. These variants were predominantly transitions (C→T, G→A), aligning with the expected alterations’ frequency of these tumor types (52) (**Fig. 2B**). As the reported calls are expected to include somatic DNA alterations, detectable also from the T/N WES analysis, and RNA-specific alterations (17), undetectable from genomics data, we quantified and described the overlap between the two call sets.

As expected from previous reports (17–19, 53), they poorly overlapped each other (**Fig. 2C**). Indeed, in both cases, variant calling was limited to a given list of target regions which, in the case of WES, is specific of the enrichment probe kit used for the sequencing experiment. Conversely, for the RNA, the set of target regions was defined using genomic coordinates of protein coding genes, as detailed in the Methods section. These differences, as expected, are not negligible, justifying on average the ∼ 41.9% of the undetected alterations from WES (**Fig. 2D**). Nonetheless, the remaining set (∼ 59.1%) of variants undetected from genomic data and identified from ENEO, despite falling within the exome capture region, are detectable only from RNA-seq (**Fig. 2D**). Conversely, we quantified and analyzed WES-variants that were not detected by our approach. The primary reason for this discrepancy is the absence of mapped reads at the genomic positions (∼ 65.9%), with a minor contribution from allele-specific lack of expression (∼ 21.6%) (**Fig. 2E**). To rule out sequencing coverage errors as the cause of the missing coverage (54), we validated both WES variants and RNA variants by comparing them with the corresponding gene expression levels. This analysis revealed significantly lower expression levels for genes harboring variants undetected by RNA-seq, compared to those detected by both methods (**Fig. 2F**, Kruskal-Wallis test, *p<* 1*e* − 5).

### The tumor-only RNA-seq approach favors the identification of immunogenic neoantigens

The simultaneous detection of germline and somatic events enables the deciphering of the alterations at the translation level, leveraging both those occurring in healthy and tumor cells. To this end, ENEO exploits the haplotype information coming from the variant calling process to phase multiple variants across the same haplotype and to generate mutated proteins (**Methods**). Using the previously described RNA-variants from TESLA patients, we generated all the possible 8-14mer mutated peptides and, coupling with patient HLA genotype, we predicted the binding likelihood as the elution rank percentile (48) using NetMHCpan 4.1 (**Methods**). As demonstrated by Reardon and colleagues (49), adopting the same threshold for elution percentile over multiple HLA to delimit the list of candidate neoantigens may result in loss of specificity or sensitivity. Thus, we computed HLA-specific optimal elution percentile thresholds over more than 100 HLA alleles using IEDB data (**Methods**) and used them to select the set of candidates pMHCs for each patient. This led to the generation of a total of 6140 pMHCs across all the cohort, resulting in 5829 candidate peptides (**Suppl. Table 2**).

We measured the sensitivity of our approach in identifying TESLA tested immunogenic peptides. Notably, applying ENEO to the tumor RNA-seq data of patients resulted in the successful identification of 26 out of the 34 positive tested immunogenic peptides, with variable performances between patients and respective histology (**Fig. 3A**). In melanoma cases (TESLA_1, TESLA_2, TESLA_3), ENEO showed the higher recall, reporting 23 out of the 26 immunogenic peptides (**Fig. 3A**). Conversely, we observed a lower resolution for the two NSCLC patients under investigation (TESLA_12, TESLA_16), where 3 out of 8 validated peptides were identified, reporting anyhow at least one immunogenic neoantigen even in the worst performing scenario. Interestingly, ENEO did not select 247 out of the total 501 (∼49.3%) of the originally predicted neoepitopes assayed as not immunogenic. This resulted in a set of neoepitopes that exhibited notable sensitivity, as indicated by the recall over increasing fractions of the proposed list (**Fig. 3B**). We compared our approach with various standard methods used by teams in the TESLA consortium, adopting the metrics defined in the original publication. These were defined as TTIF (Top 20 Immunogenic Fraction) and FR (Fraction Ranked) and are used for comparisons within individual patients. The first is a proxy of the specificity, as it approximates the number of immunogenic peptides in the first 20th positions of the ranked list, considering 20 as an adequate testing size for therapeutic application (22). The second could be intended as a sensitivity metric, reporting the number of immunogenic peptides over the total positive tested in the first 100th positions of the ranked list. All the participant teams used T/N WES for variant calling and, consequently, the list of experimentally tested peptides is expected to come from variants detectable from WES. ENEO outperformed on average other teams for all melanoma patients in both FR and TTIF (one-sided t-test, *p<* 1*e* − 4; **Fig. S3**). For NSCLC patients, ENEO demonstrated superior FR for patient TESLA_16 than average (one-sided t-test, *p <* 1*e* − 4; **Fig. S3**), while for patient TESLA_12, the result was not significantly different from other teams’ submissions on average (two-sided t-test, *p* = 0.5; **Fig. S3**). Of note, in 3 patients we identified a previously characterized cancer neoantigen (RVWDVSGLRK) with strong predicted binding affinity, known to come from an A-to-I RNA editing event occurring in the protein COPA in the 164^th^ aminoacid, responsible for a substitution of an isoleucine with a valine. This event was recently proposed as a driver of metastasis in colorectal cancer through ER stress (55), and was eluted in melanoma samples following MHC-purified mass spectrometry and demonstrated to induce IFN_*γ*_ secretion (56, 57).

**Fig. 3.**
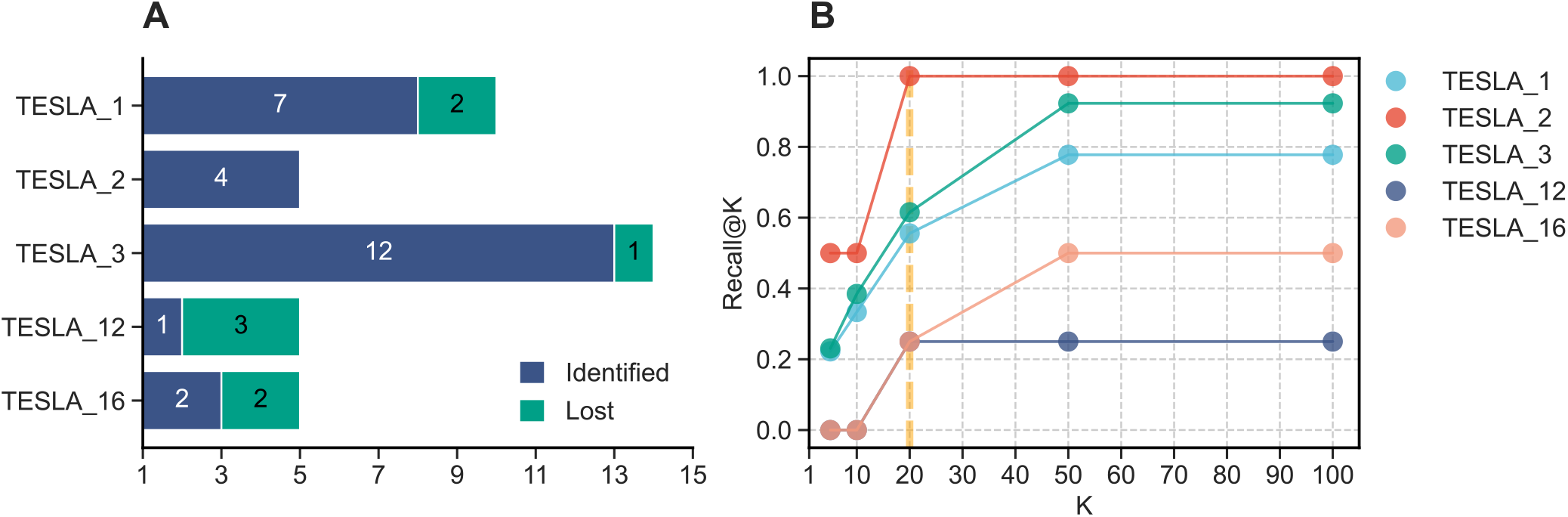
Performances on immunogenic pMHC identification from the TESLA benchmark. **A**. Number of positive experimentally validated pMHC detected or non-detected by ENEO predictions. **B**. Patient-wise Recall-at-K (Recall@K), defined as the fraction of immunogenic pMHCs identified for each patient (Recall) over increasing fractions (k) of the proposed ranked list. The scattered orange line is fixed at the 20th position, preserving the choice adopted in the original TESLA publication.

### Neoantigen prediction is consistent across different tumor types and mutational burden levels

To verify the robustness of our approach, we collected tumor RNA-seq from two publicly available cohorts, that applied a similar screening protocol in two different tumor types. The first one, here referred to as “Gros”, examined the antigen specificity of T-cell responses detected from peripheral blood lymphocytes in three metastatic melanoma patients (23). The second one, here referred to as “NCI”, is composed of patient with metastatic gastrointestinal cancers characterized by microsatellite stability and mismatch repair proficiency (5, 6). In this group, neoantigen specificity was determined from the T-cell response of cultured tumor infiltrating lymphocytes. These two studies shared the screening protocol, consisting in somatic variant calling from T/N WES and, after transcript expression confirmation and patient-specific variant selection, screening via synthesis of 25-mer constructs and transfection in autologous dendritic cells using tandem mini genes (58).

For the Gros cohort, we processed tumor RNA-seq reads with ENEO and merged the resulting candidate pMHCs into 25mer frames, resulting in the production of 612 constructs (Table xxx). Our analysis identified 31, 32, and 30 out of all tested 25-mers, successfully detecting 6 out of the 7 positively tested 25-mers. Furthermore, we examined the minimal epitope immunogenicity of positively tested 25-mers to determine how many positive pMHCs were reported by ENEO and their relative rankings. We identified 8 out of 9 positive pMHCs, with at least one immunogenic peptide detected per patient (**Fig. 4A**). Importantly, among the tested constructs, immunogenic neoantigens were placed by ENEO within the top 30 ranked pMHCs (**Fig. 4B**). Owing to the patient-specific ranked list, all the positive tested pMHCs were at least included in the upper half (i.e., 50^th^ percentile) of the candidates (**Fig. 4B**). Notably, the peptide FVVPYMIYLL, immunogenic for patient Mel_3998, derives from two phased somatic variants (p.Y1000F and p.H1007Y) in the PDS5A gene. Its detection was possible due to the retention of haplotype information by the variant calling process and its use in subsequent variant peptide generation (**Methods**).

**Fig. 4.**
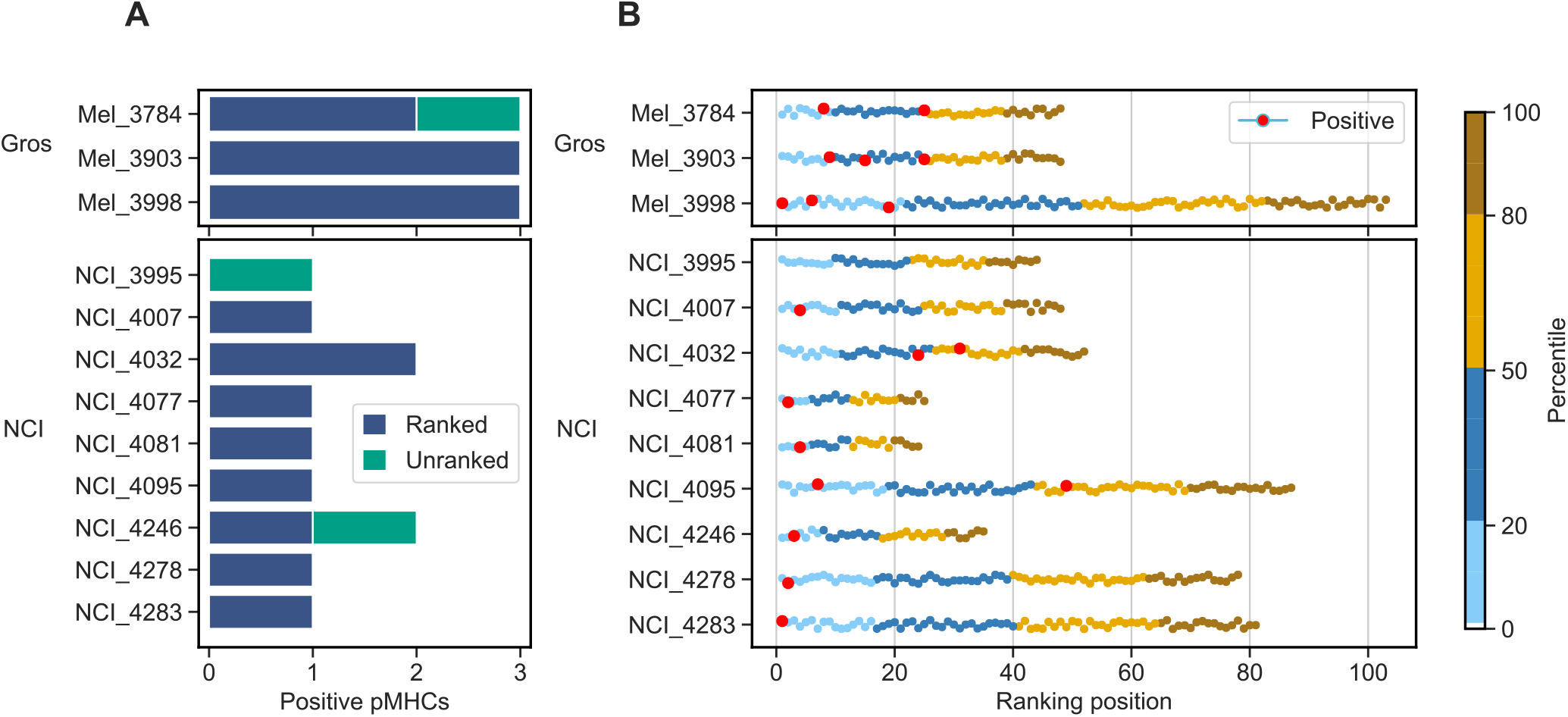
Identification of pMHCs in external cohorts. **A**. Number of positive pMHCs identified by ENEO for each patient. **B**. Patient-wise ranking of candidate pMHCs for tested 25-mer identified by ENEO, colored by their percentile among the ranked list.

Given the availability of somatic variants profiled by authors via T/N WES, we could investigate origin of the missed neoantigen. The variant underlying the peptide KVDPIGHVY, responsible for the p.E168K transition in

MAGEA6 (a known melanoma-associated antigen (59)), was discarded due to insufficient coverage (*<* 5 RNA-seq reads). This is in accordance with Gartner et al.’s findings (24), which reported no distinct evidence for the alternative allele in the transcriptomics data. As the experimental procedure followed by authors was developed on WES variants, RNA-exclusive variants detected by our approach were not subject to the original immunological screening. We reported an average of 171 25-mer constructs generated from just as many RNA-exclusive variants, which leads to the proposal of ∼319 candidate pMHCs for each patient (**Suppl. Table 3**).

For the “NCI” cohort, we resembled the experimental procedure adopted for the “Gros” one. This group of patients represents an even more challenging experimental setup for testing the robustness of ENEO, as these tumors have low burden of somatic mutations and are less responsive to checkpoint inhibitor therapy (60). Tumor RNA-seq analysis led to an average of ∼271 constructs, with an extended difference between patients (**Suppl. Table 3**). We successfully identified 11 out of the total 13 positively tested 25-mer, detecting at least one immunogenic construct for all but one patient (**Fig. 4A**). In the ranked list of pMHCs generated out of the tested 25mers, immunogenic neoantigens were found within the top 10 ranked entries, corresponding to the first 20th percentile, for 7 out of the 9 patients under investigation (**Fig. 4B**). ENEO remained robust to the heterogeneity of this dataset, in term of sequencing platform, read length and, most importantly, to the overall number of reads (**Suppl. Table 4**). As even for this cohort somatic variants profiled via T/N WES are available, we investigate events behind missing neoantigens. For the patient NCI_3995, the p.G12D transition in KRAS detected from T/N WES is reported as the source for the single immunogenic peptide reported. In the RNA-seq data we failed to observe consistent evidence for the mutated transcript, as the mutated allele is supported by only 2 reads out of the total 8 mapped in this region. A similar scenario is reported for the patient NCI_4246, where the unidentified peptide arises from a SNV in the gene ARMC9, encoding for a melanocyte-specific antigen (KU-MEL1). For this variant, we found that in the RNA-seq data the mutated allele is supported only by two out of the total 6 reads mapped. In both cases, the alteration is filtered out during the variant calling process due to the low coverage, resulting in a poor genotype quality. For this group of patients, as for the Gros cohort, the analysis of RNA-seq data leads to the proposal of 264 untested candidate 25-mers, originating from the same number of RNA-specific variants. The binding affinity estimation and subsequent filtering of ENEO results in the generation of ∼1150 candidate pMHCs (**Suppl. Table 3**).

Taken together, these results indicate that the tumor-only approach implemented in ENEO permits the identification of cancer neoantigens in different histological and technical conditions, without requiring neither a matched control nor concurrent DNA-sequencing.

## Discussion

In this study, we investigated the feasibility and limitations of predicting immunogenic neoantigens using only the tumor RNA-seq data. We compared our approach with state-of-art methods that additionally require the tumor/normal (T/N) DNA sequencing on a benchmark dataset, and validated it on external cohorts encompassing different tumor histologies and experimental setups. We accomplished this by developing ENEO, an easy-to-use and automatized computational workflow designed to predict tumor neoantigens using RNAseq data without a matched-control sample. Being built with the Snakemake workflow, it is designed to be modular, extensible and scalable to different platforms and infrastructures. Importantly, ENEO addresses the challenge to distinguish germline variants in the absence of a matched control sample through a Bayesian probabilistic model. The model leverages variant population allele frequencies and genotype likelihood and proves to be able to remove most germline variants out of the detected RNA-variants in the TESLA cohort. We tested its ability to detect immunogenic neoantigens in public studies, encompassing different tumor types and experimental validation, proving its clinical utility even in very low mutated tumors like MMR-proficient gastrointestinal cancers. Harnessing the growing evidence pointing to the contribution of transcriptional specific alterations to the plethora of MHC-presented peptides, we demonstrated that a tumor RNA-based approach like the one implemented in ENEO preserves resolution on canonical mutated peptides originated from expressed genomic alterations, proposing at the meanwhile candidate pMHCs specifically arising from RNA-specific mutations. Despite the encouraging performances, the reported approach is not exempt of limitations. Due to the nature of an RNA-seq experiment, the resulting reads are limited to the actively transcribed regions and proportional to the amount of sampled mRNA molecules and to their length. In a typical short read RNA-seq experiment, the number of produced reads is often set to a desirable threshold, which is defined *ad hoc* to satisfy study requirements. This introduces a significant limitation in the resolution of the generated experiment which, although being still adequate for a general gene expression analysis, will not confer enough power to the detection of a large repertoire (61) of somatic mutations. Indeed, in our tests/experiments, we consistently observed that missed neoepitopes are generated from variants falling in under-represented regions, where the minimum coverage requirement was not met, causing uncertainty in the genotyping process. As most of the tested datasets were generated with a relatively low depth protocol (*<*50M reads), recalling previous reports (61), we speculated that increasing the number of sequencing reads could dramatically boost the sensitivity of an RNA-seq based approach, enabling the detection of tumor specific alterations even in low expressed transcripts.

Another important consideration must be accounted due to the lack of a matched control sample. While we reported that the use of population genetic databases and genotype likelihoods via the Bayesian germline error model of ENEO is fruitful, rare and ultrarare events may be still prone to misclassification. Additionally, population databases are known to be still under-representative of many ethnicities, suggesting that this may negatively affect the performances on them. These limitations can potentially be mitigated using larger population databases, which, at time of writing, are becoming increasingly representative of previously poor characterized ethnicities (62).

Overall, ENEO opens to the applicability of neoantigen detection using solely the tumor RNA-seq data, providing a faster and cost-effective way to interrogate the transcribed regions of the human genome. Its robustness across different tumor histologies and mutational burdens highlights its potential in various translational and experimental setups, reinforcing the importance of high-resolution transcriptional profiling in the precision oncology.

## ACKNOWLEDGEMENTS

We acknowledge the CINECA award under the ISCRA initiative, for the availability of high-performance computing resources and support.

## AUTHOR CONTRIBUTIONS

DT conceived and conducted the study, developed methods, performed analyses, interpreted the results, and drafted the manuscript; MD contributed to the initial conceptualization, methods development and methodology; GB formalized and developed the statistical model used in the study; GG and MC participated in study conception; RD supervised the study, participated in study conception, data interpretation, and drafted the manuscript. All authors reviewed and approved the final manuscript.

## Supporting Information for

**Fig. S1.**
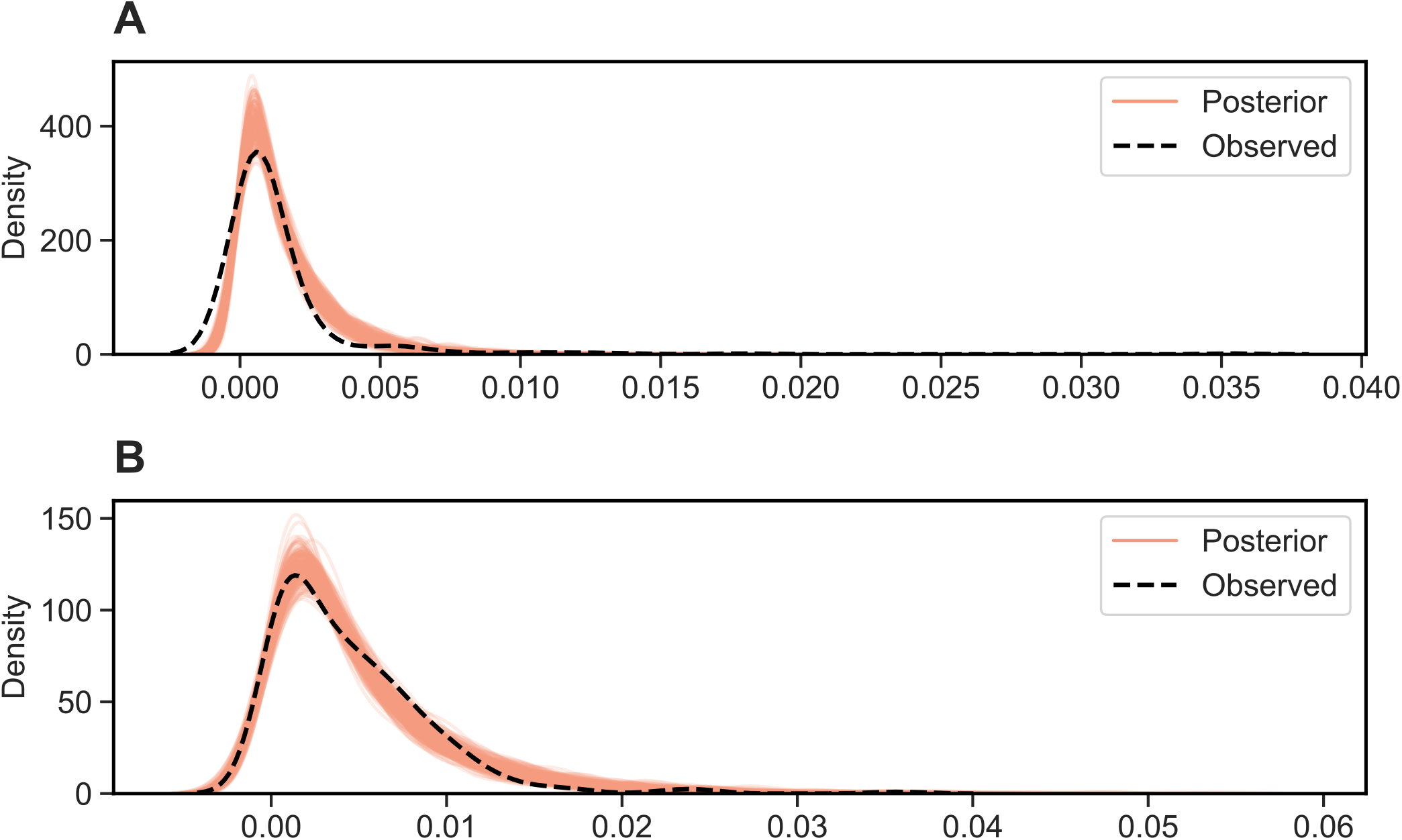
Posterior density plot for parameters estimated with Metropolis Hastings using 200 values for alpha and beta from the HPD. A. Posterior estimates for the exon mutation rate compared to observed data. B. Posterior estimates for the base mutation rate compared to observed data.

**Fig. S2.**
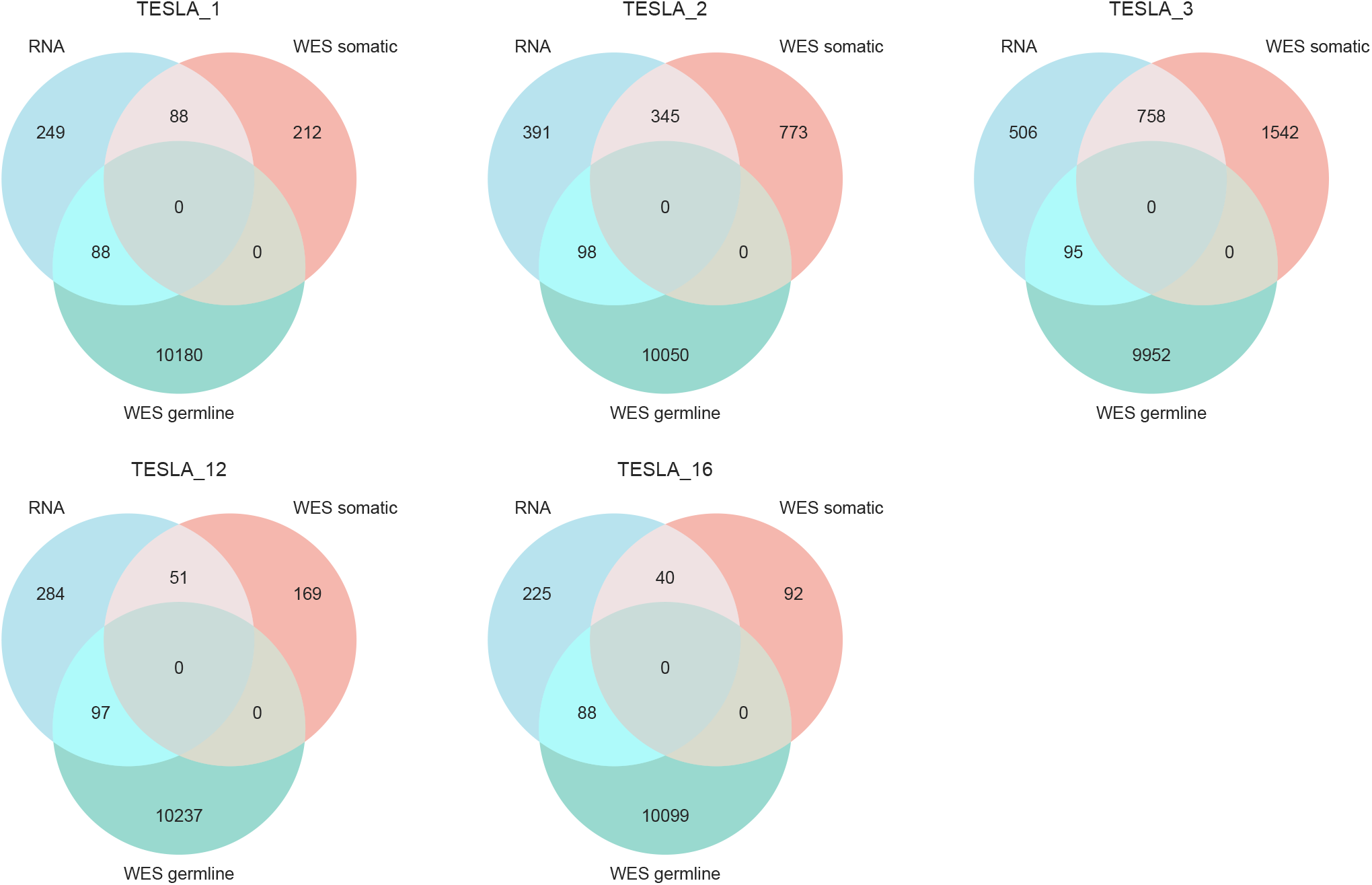
Patient-wise measured overlap between variants reported to have an impact on the downstream protein sequence for RNA, WES somatic and WES germline, after filtering RNA-variants to retain only those with a computed somatic probability above 0.5.

**Fig. S3.**
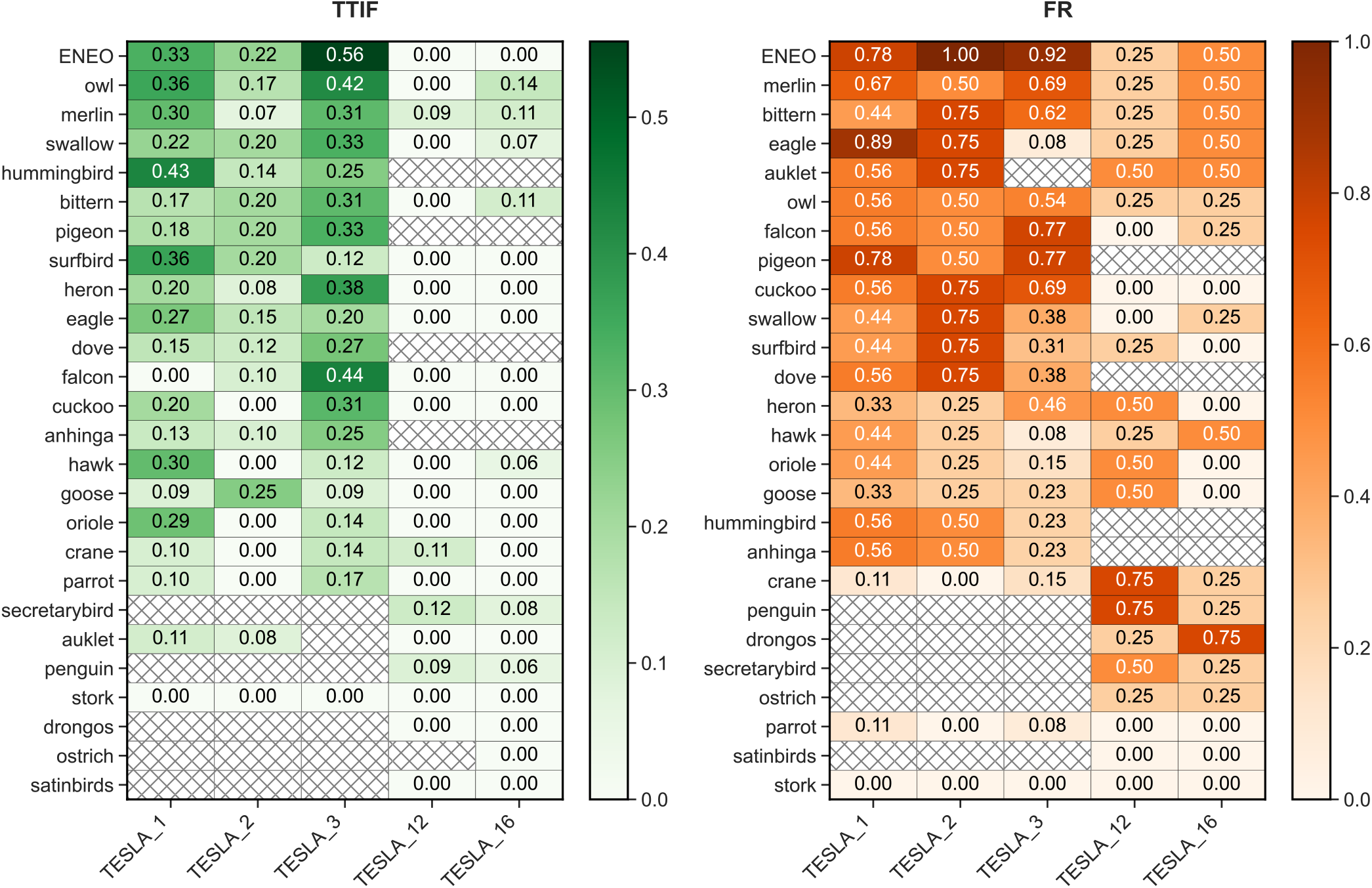
Patient wise comparison between ENEO and other TESLA teams on neoantigen predictions using metrics from the original publication. TTIF (Top 20 Immunogenic Fraction) refers to the number of immunogenic peptides in the first 20^th^ position of the ranked list relative to the tested peptides. FR (Fraction Ranked) measures the number of detected immunogenic peptides over the total positive tested peptides in the first 100^th^ positions of the ranked list. A crossed box represents a missing submission.

